# Assessing State-Specific Accuracy of Cofolding Models for Kinases and GPCRs

**DOI:** 10.64898/2026.05.06.723350

**Authors:** Leon Obendorf, Niklas Piet Doering, Petra Knaus, Gerhard Wolber

**Affiliations:** Department of Biology, Chemistry and Pharmacy, Institute of Biochemistry, Signal Transduction Group, Freie Universität Berlin, Thielallee 64, 14195 Berlin, Germany; Department of Biology, Chemistry and Pharmacy, Institute of Pharmacy, Molecular Design Group, Freie Universität Berlin, Königin-Luise-Str. 2+4, 14195 Berlin, Germany

## Abstract

AI-driven cofolding models have emerged as powerful tools for predicting protein–ligand complexes, yet whether ligand placement faithfully captures the conformational states of dynamic proteins remains unclear. Here we show that cofolding adaptively remodels binding pockets around bound ligands, but that this local accuracy is frequently decoupled from recovery of the broader conformational state. We benchmark four models, AlphaFold3, RosettaFold3, Boltz-2, and Chai-1, against a set of kinases and class A G protein-coupled receptors (GPCRs), protein families whose pharmacology depends on well-defined structural states. We find that even when ligand root-mean-square deviation (RMSD) is low, critical state markers, including kinase activation-loop geometries and GPCR intracellular arrangements, are frequently mispredicted. Incorporating state-annotated templates and filtered multiple sequence alignments (MSAs) improves conformational recovery in selected cases, yet weakly impacts others. Furthermore, while orthosteric ligand placement is generally reliable, allosteric binders expose a consistent blind spot across all models. These findings establish conformational decoupling as a fundamental limitation of current cofolding approaches, with direct implications for state-selective drug design.

## Introduction

Proteins act as dynamic macromolecules that reside in an ever changing conformational ensemble.^1^ This ensemble often determines physiological outcomes and can be modified by ligand-binding events. Kinases^2^ and G protein-coupled receptors (GPCRs)^3^ are two well known protein families where these shifts in conformational ensembles guide activity and downstream signaling. Although kinases share a conserved bilobal fold, they differ in how active and inactive states are achieved, often through small shifts in the activation loop, the DFG motif, the *α*C-helix or nearby regulatory elements that affect substrate binding.^4^ In GPCRs, signaling arises from rearrangements of conserved motifs (e.g. DRY, NPxxY, PIF) and a reshaping of the intracellular binding pocket to open for transducer binding.^3^ When aiming to work with structural data of kinases or GPCRs in *in silico* investigations, it is crucial to model the correct state for the bound ligands.

Although deep learning models have transformed structural biology through *de novo* protein structure generation (protein folding), widely used predictors such as AlphaFold2 (AF2) often default to the most common conformations present in their training sets.^5^ For systems that adapt significantly different states, this can lead to problems such as prediction in an unwanted state or even prediction in mixed states that represent an non-physiological average of all states within the training set.^5^

Methodological advances aim to address state ambiguity in deep-learning structure prediction by biasing protein prediction models toward user-specified functional conformations. One widely used strategy is template filtering, in which inputs are restricted to state-annotated experimental structures from curated resources such as GPCRdb (https://gpcrdb.org/)^6^ and KLIFS (https://klifs.net/),^7^ enabling predictions to be steered toward desired conformers.^8^ A complementary approach modulates the evolutionary signal contained in multiple sequence alignments (MSAs): reducing MSA depth or clustering sequences can weaken dominant co-evolutionary patterns and expose alternative structural states, including inactive and DFG-out kinase conformations^9,10^ and alternative states of GPCRs.^11^ Importantly, these two strategies are not independent. Heo and Feig demonstrated that state-annotated templates alone are insufficient when deep MSAs are present, whereas no or shallow MSA sampling allows templates to reliably enforce active and inactive GPCR states with near-experimental accuracy.^5^ This combination was also shown for kinases to outperform template-only input, particularly in flexible or poorly conserved regions.^8^ Together, these studies show that state-aware prediction is achievable, but remains highly sensitive to careful manual template selection combined with MSA control rather than emerging robustly from the models themselves.

In recent developments, cofolding of small molecules with proteins was made possible by AlphaFold-3 (AF3)^12^ and soon Boltz-2,^13,14^ Chai-1^15^ and recently RosettaFold-3 (RF3)^16^ followed. Trained on diverse biomolecular interactions, they are supposed to be able to geometrically model induced fit and extend the rigid-body assumptions of conventional docking. In benchmarks such as “PoseBusters”,^17^ these models achieved success rates that surpass widely used docking tools like AutoDock Vina^18^ and GOLD^19^.^20^ However, these gains primarily reflect improved geometric placement rather than explicit learning of physical interaction energetics. As shown by Masters et al., ^21^ systematic perturbation of binding sites often failed to alter predicted ligand poses, with models placing ligands in canonical positions even after mutations abolished the pocket. Subsequently, this motivates testing whether state-biasing strategies developed for classical protein folding, such as MSA manipulation and template selection, improve protein–ligand cofolding and state-specific binding-site geometry.

We systematically assessed the practical capabilities and limitations of current protein-ligand cofolding approaches in state-dependent systems using four state-of-the-art models: AF3,^12^ Boltz-2,^13,14^ Chai-1,^15^ and RF3.^16^ We curated a benchmark of kinases and Class A GPCRs spanning distinct functional states absent from all model training sets, enabling prospective rather than retrospective evaluation. Controlled template and MSA biasing strategies, informed by state annotations from KLIFS^7^ and GPCRdb,^6^ were applied throughout. For kinases, we selected structures that were recently solved and comprise all three inhibitor types: LRRK2 bound by the type II inhibitor RN277 (PDB: 9DMI),^22^ the PI3K^H1047R^ mutant bound by the type III inhibitor Zovegalisib within a cryptic pocket (PDB: 8TSD),^23^ and the active-like state of Activin-like kinase ALK2 kinase bound by the type I inhibitor RK783 (PDB: 9L04),^24^ supplemented by an AMPPNP-bound ALK2 structure to probe nucleotide recognition beyond inhibitor-induced states (PDB: 6UNQ).^25^ The GPCR subset comprised three Class A receptors spanning inactive and active signaling states: the antagonist-bound 5-HT_2A_ receptor, the agonist-bound adenosine A_2A_ receptor, and the δ-opioid receptor engaged by a G protein-biased agonist, systems that require simultaneous recovery of orthosteric ligand placement and the intracellular geometries associated with receptor activation, providing a stringent test of whether ligand guidance alone is sufficient to enforce global conformational states.

## Results

### Ligand placement is robust across cofolding models but decoupled from state accuracy and pocket geometry

We first evaluated ligand placement and binding-site geometry in complexes predicted by the cofolding frameworks. Across the evaluated kinase systems, low ligand RMSD did not consistently coincide with correct recovery of binding-site geometry. The first example showing this pattern is LRRK2 bound to the type II inhibitor RN277. Here, across predictions that reproduced the global kinase fold, ligands were positioned consistently (Figure 1A) and ligand RMSDs were general below 0.8 Å. Regarding binding-site geometry, the LRRK2-RN277 reference PDB structure features an upward-oriented conformation of the DYG-motif tyrosine, a geometry not observed in PDB entries released before the 2025 cryo-EM structure. This novel Y2018 orientation (Figure 1B) was recovered only in a subset of predictions: by AlphaFold3 with biased inputs and in some Chai-1 predictions. Boltz-2 and RosettaFold-3 instead consistently reproduced the canonical inactive tyrosine arrangement. By simple RMSD calculation, Chai-1 using the `no-msa custom templates` condition yielded the lowest ligand RMSD (0.42 Å), despite wrong Y2018 geometry and a non-physical chair deformation of a phenyl moiety in RN277 (Figure 1C), with the latter being present in all Chai-1 predictions.

**Figure 1:**
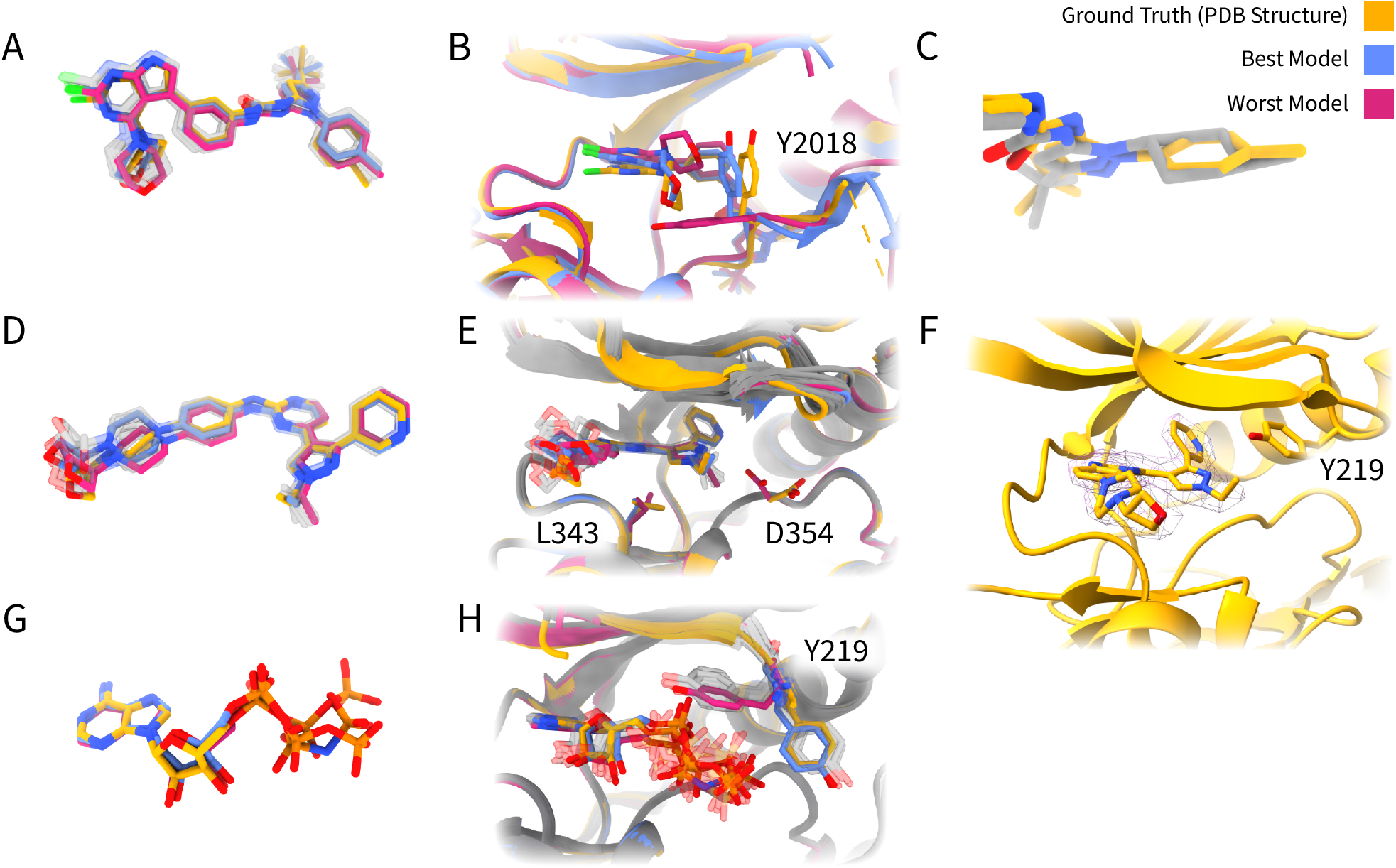
Cofolding predictions of kinases with small molecule inhibitors compared to groundtruth structures: **A-C** LRRK2 with the type II inhibitor RN277, **D-F** ALK2 with the type I inhibitor RK783, **G-H** ALK2 with ATP bound. **A** Superposition of all predicted RN277 ligand conformations (transparent) with the cryo-EM reference structure (PDB: 9DMI; yellow), highlighting the structurally closest (red) and farthest (blue) structure. **B** Close-up view of the DYG-motif with Y2018 shown in stick representation in the predicted structures compared to the reference. **C** Comparison of groudtruth (yellow) with Chai-1 predicted ring conformation of RN277. **D** Superposition of predicted RK783 ligand conformations in ALK2 with the crystal structure reference (PDB: 9L04; yellow), showing differences in to solvent-exposed distal portion of the ligand (lower left) and orientation of an ethyl group. **E** Close-up view of the RK783 binding site with deviations in local pocket geometry in the worst prediction where D354 is rotated toward the ligand, and the 1-ethyl subsituent of a pyrazole is facing the lower kinase lobe. **F** Orientation of the experimentally determined 1-ethyl substituent of RK783 relative to the p-loop and Y219 in the groundtruth PDB: 9L04. **G** Superposition of best and worst predicted ATP conformations with the AMPPNP-bound reference structure (PDB: 6UNQ). **H** Close-up view of the nucleotide binding pocket showing the conformation of the P-loop residue Y219 relative to the bound nucleotide. The outward-facing Y219 observed in the reference structure is reproduced in a subset of predictions.

Similar for ALK2 bound to the type I inhibitor RK783, models successfully predicting the kinase fold generated also ligand RMSDs below 0.8 Å. Structural deviations in the lig-and were localized primarily to solvent-exposed ligand regions and adjacent pocket residues (Figure 1D). Furthermore the 1-ethyl substituent of RK783 pointing toward Y219 was recovered in a subset of predictions. Among the successful predictions, AlphaFold3 with biased MSAs shows the lowest ligand RMSD (0.57 Å). The Boltz-2 condition with `biased MSA + no templates` reproduced the ethyl orientation but showed a 180° flip of the pyridine ring in RK783. Chai-1 under the `no MSA + no templates` condition resulted in incorrect ethyl orientation and rotation of D354 toward the ligand, disrupting a salt bridge observed in the experimental structure between D354–R375 (Figure 1E). The variation in ethyl-group orientation in the predictions contrasts with the experimental density map, which indicates a defined substituent position (Figure 1F).

To extend the analysis beyond inhibitors, ATP binding to ALK2 was evaluated using the AMPPNP-bound structure (PDB: 6UNQ) as a reference. Based on its deposition date, 2019, this structure was likely present in all model training datasets. Ligand RMSDs ranged from 0.58 to 1.40 Å across successful predictions, with protein backbone RMSDs below 0.5 Å. A defining feature of the active site is the outward rotation of the P-loop residue Y219, which was reproduced only in a subset of predictions (Figure 1H), from which none was by Chai-1. Boltz-2 achieved the lowest ligand RMSD (0.58 Å) using standard MSA generation without templates.

Across all three systems, low ligand RMSDs were also observed in models that differed in residue orientation and local interaction networks, indicating that accurate ligand placement can occur without recovery of the experimental binding-site geometry.

### Allosteric binding pockets are not sampled by cofolding

To evaluate whether cofolding can recover non-orthosteric binding modes, we analyzed the PI3K*α*-H1047R mutant in complex with Zovegalisib (RLY-2608), a first-in-class, mutant-selective allosteric inhibitor.^23^ In the experimentally determined structure (PDB 8TSD), Zovegalisib occupies a cryptic allosteric pocket distinct from the canonical ATP-binding site. However, across all models and parameter settings tested, none of the predicted structures placed the ligand in this allosteric pocket; instead, every model positioned it within the ATP binding site, as shown in Figure 3 2.

**Figure 2:**
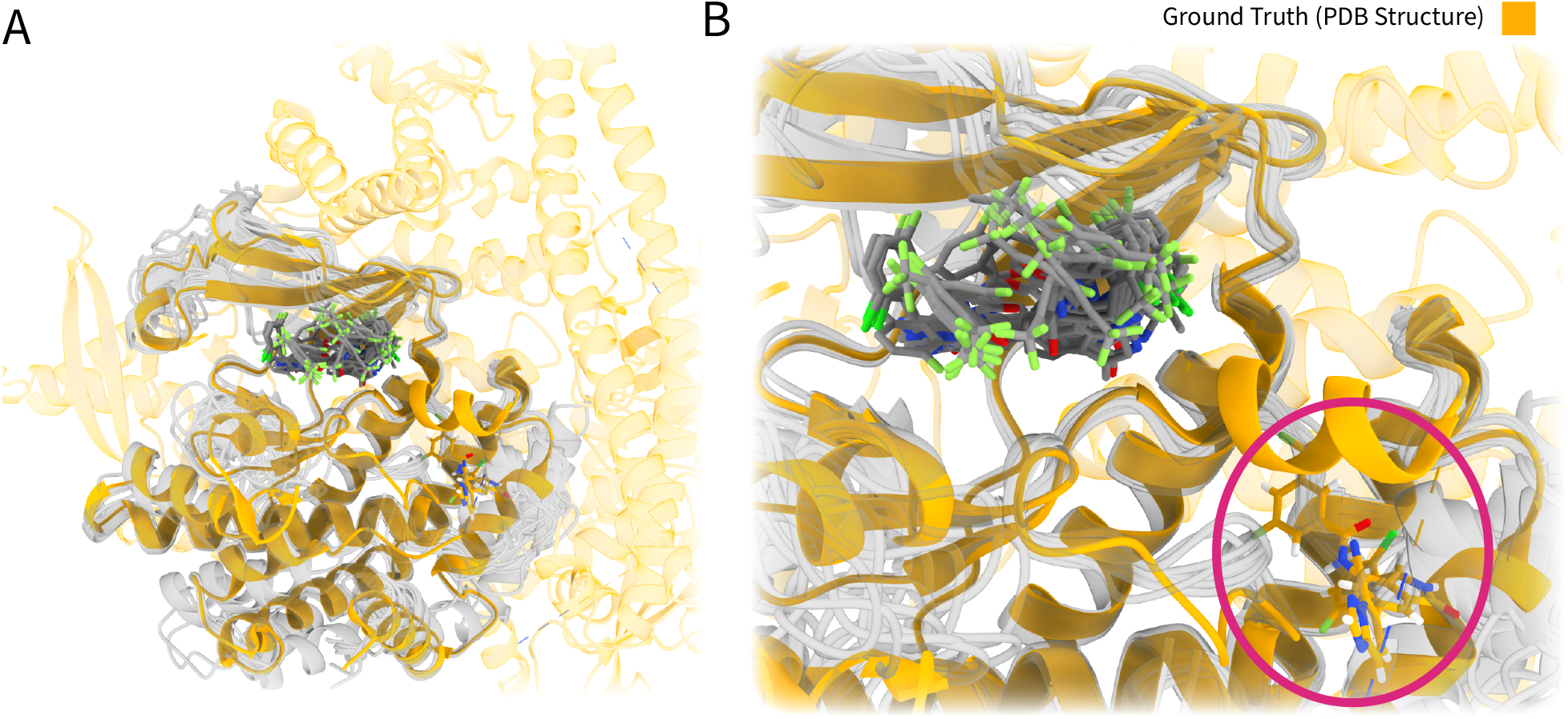
PI3K*α*-H1047R mutant in complex with Zovegalisib (RLY-2608). Yellow indicates the ground-truth PDB structure, while grey shows all cofolding model predictions. **A** Overview of the general PI3K*α* fold. **B** Ligand placement of Zovegalisib (RLY-2608). All cofolding models position the ligand in the canonical ATP-binding site, whereas the true ligand occupies a nearby allosteric pocket (red circle).

**Figure 3:**
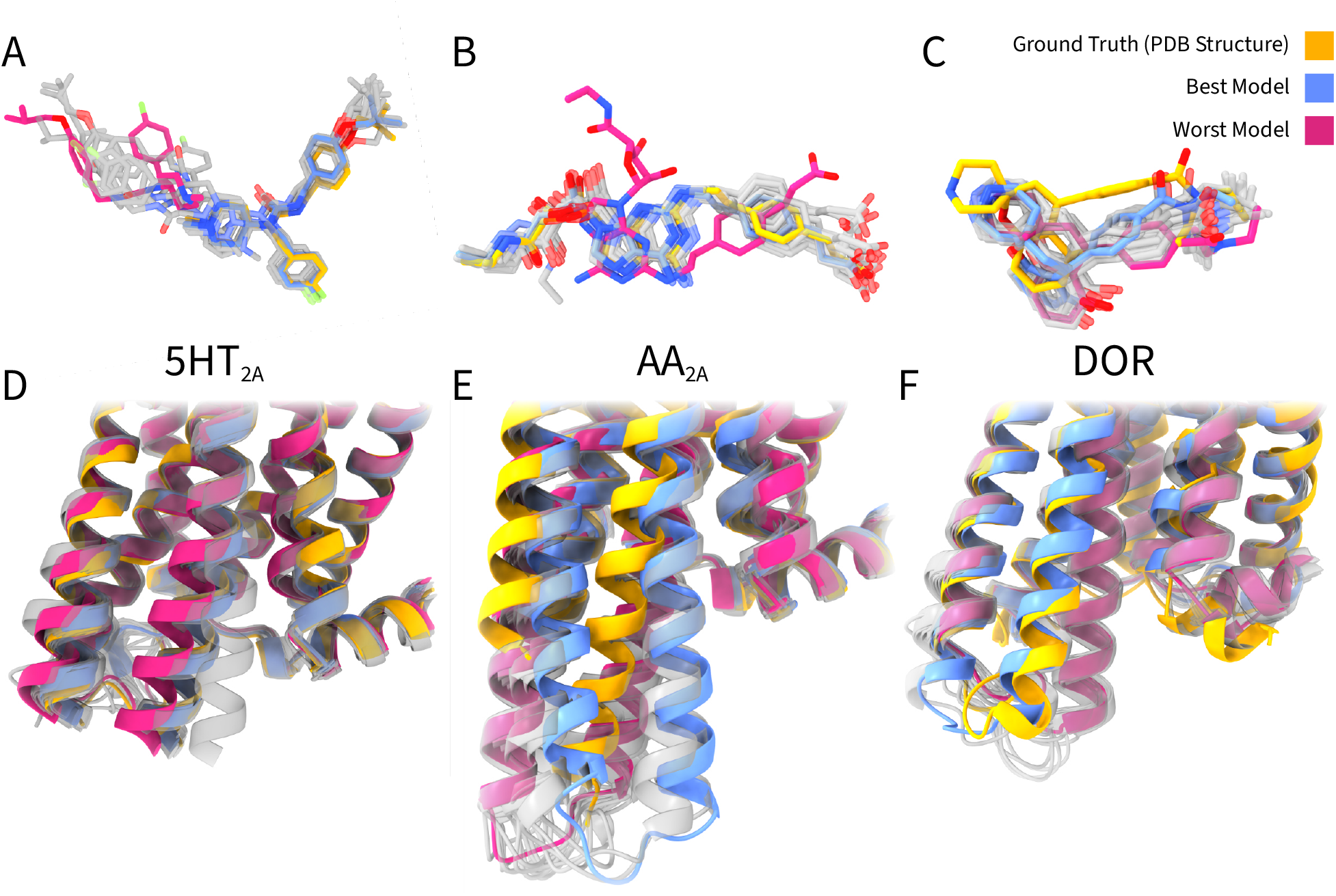
Placement of ligands and ICL3s in the cofolded GPCR structures. Yellow indicates the reference PDB (ground truth), blue shows the best-performing model based on ligand RMSD, red shows the worst-performing model, and all other predictions are represented in grey. **A** Conformations of the antagonist pimavanserin in the HT_2A_ receptor. **B** Conformations of the agonist CGS21680 in the adenosine A_2A_ receptor. **C** Conformations of the G protein-biased agonist ADL5859 in the DOR. **D** ICL3 folds of the inactive-state HT_2A_ receptor models. **E** ICL3 folds of the active-state A_2A_ receptor models. **F** ICL3 folds of the active-state DOR models.

Despite the incorrect ligand placement, the overall kinase fold was reproduced accurately, and backbone RMSDs were generally below 1.5 Å. Efforts to guide sampling, including blocking the ATP site sterically, introducing known protein interaction partners associated with the allosteric conformation, and using Boltz-2 binding contact flags to bias the ligand to-ward the cryptic pocket, did not lead to recovery of the allosteric site. All models continued to place the ligand in the canonical ATP pocket. These observations indicate that while cofolding reliably reproduces global protein architecture, it does not spontaneously sample cryptic or allosteric binding modes, even when guided. This highlights a limitation of current cofolding approaches for predicting ligands that occupy non-canonical sites, even when the overall fold is accurately modeled.

### GPCR ligand placement is decoupled from intracellular state recovery

To assess the ability of cofolding models in capturing both ligand binding and receptor conformational changes, we examined three Class A GPCRs representing distinct active and inactive states. GPCRs undergo substantial rearrangements, particularly in intracellular regions such as ICL3, which play key roles in G protein coupling. We selected three representative test cases, the inactive-state 5-HT_2A_ receptor bound to pimavanserin (PDB: 8ZMG)^26^ (Figure 3 A,D), the active-state adenosine A_2A_ receptor bound to CGS21680 (PDB: 8UGW)^27^ (Figure 3 B,E), and the active-state δ-opioid receptor (DOR) bound to the G protein-biased agonist ADL5859 (PDB: 8Y45) ^28^ (Figure 3 C,F). Across these examples, a consistent pattern emerged: while ligand placement and pocket adaptation within the orthosteric binding site could often be predicted accurately, the intracellular regions, particularly ICL3, were frequently misfolded, revealing a decoupling between ligand binding and intracellular state recovery.

Across the three GPCRs, ligand placement was generally accurate, with all models not only correctly positioning the ligand within the orthosteric binding site but also, in most cases, reproducing the correct ligand conformation (Figure 3 A-C). For the adenosine A_2A_ receptor, nearly all models produced reasonable ligand placements, with the exception of Chai-1 when using shallow MSAs, which led to less accurate positioning. A notable exception among the predictions were all Chai-1 outputs in the inactive-state 5-HT_2A_ receptor bound to pimavanserin, where only the inclusion of structural templates allowed recovery of the correct ligand pose and conformation (Figure 3 A). In the δ-opioid receptor bound to the G protein-biased agonist ADL5859, all models showed some deviations in ligand positioning, yet the general placement and overall ligand geometry within the binding pocket remained correct (Figure 3 C). These observations indicate that, despite occasional outliers, ligand prediction is often robust across both active and inactive GPCR states.

In contrast to ligand placement, the accuracy of the protein fold varied considerably across the receptors. While residues within the binding pocket were generally well-modeled, larger discrepancies became apparent when examining the intracellular regions, such as the ICL3, where conformational differences between active and inactive states are most pronounced (Figure 3 D-F). The inactive-state 5-HT_2A_ receptor was folded relatively well across models (Figure 3 D), but both active-state receptors, the adenosine A_2A_ receptor and the DOR, were modeled suboptimal (Figure 3 E-F). This was particularly evident in ICL3, where nearly all models failed to recover the correct active-state conformation. Often, ICL3 adopted an inactive-like fold or a “hybrid” configuration. In the DOR model we also observed that the ICL3 geometry appears to influence ligand placement, with displacements of ICL3 leading to corresponding shifts in the binding pocket (Figure 3 C,F). The only notable exception to ICL3 misfolding was the DOR model generated with AF3 using custom MSAs and templates, which closely approached the correct active-state intracellular geometry. These results highlight a clear divergence between robust ligand prediction and the challenges of accurately modeling the actual protein fold, particularly in active-state GPCRs.

Overall, these results underscore a decoupling between ligand placement and intracellular state recovery. While current cofolding models can often position ligands correctly within the orthosteric site, they struggle to reproduce the precise intracellular conformations associated with a specific activation state. This has important implications for GPCR modeling, suggesting that cofolding may be useful for guiding ligand-based decisions, as ligand placement can often be achieved even when intracellular loops are mispredicted. However, it does not allow one to reliably infer the functional outcome of a ligand, since accurate modeling of the intracellular signaling interfaces is still largely beyond reach without additional structural guidance.

### Template and MSA biasing affect prediction quality but do not reliably enforce correct conformations

The importance of the presence of either MSA or of biased templates becomes clear across systems as omitting MSAs and templates in AlphaFold3 and RosettaFold-3 results in globally misfolded structures with backbone RMSD values exceeding 17 Å and misplaced ligands (Figure 4A). Under the same conditions, Chai-1 and Boltz continued to generate overall correct protein folds without the use of MSAs or templates, but frequently exhibited substantial errors in ligand conformation or pocket geometry. In two cases, these issues produced the poorest-quality structures among all non-globally misfolded predictions.

**Figure 4:**
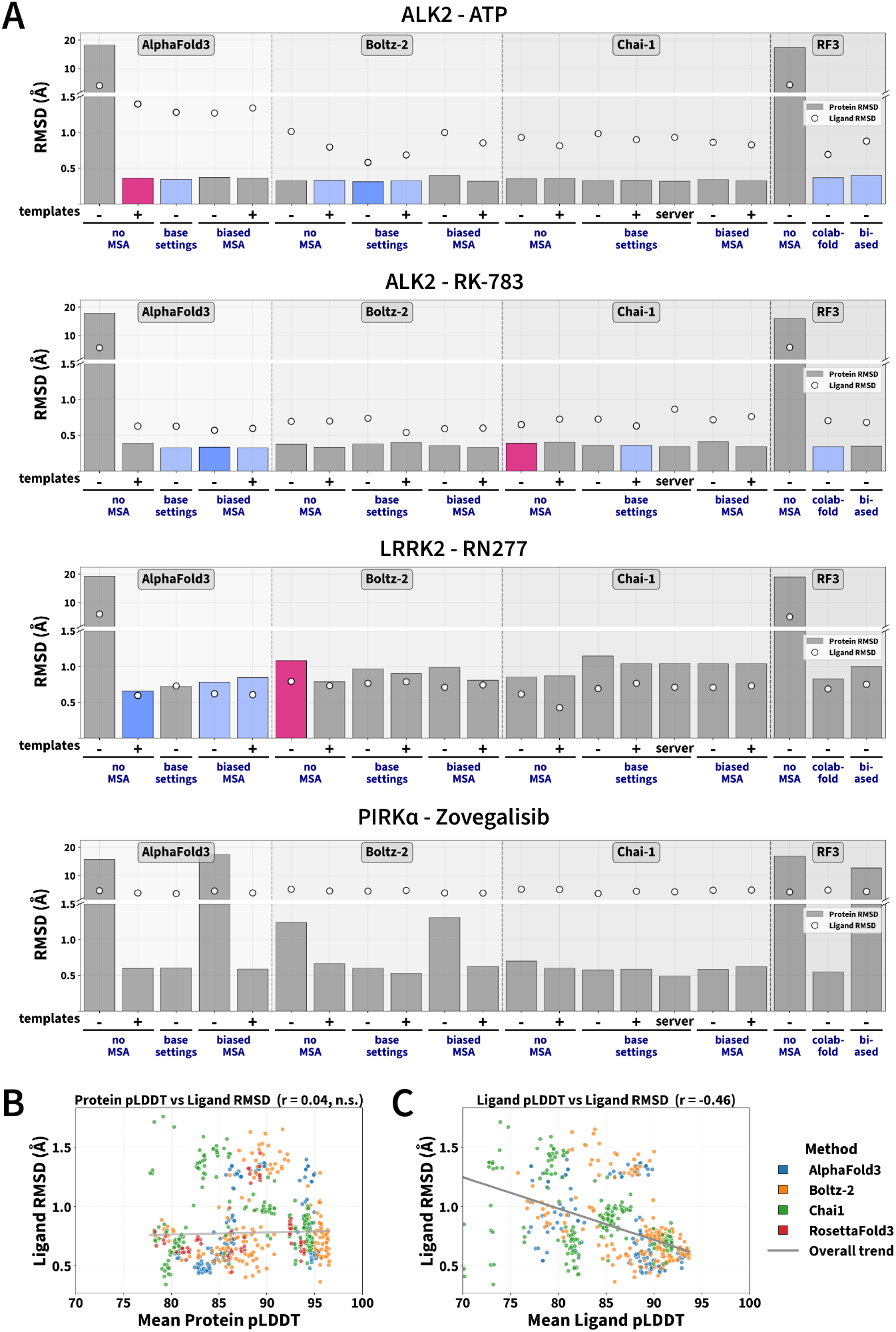
**A** Protein backbone RMSD (bars) and ligand RMSD (dots) for the lowest Protein pLDDt cofolding predictions generated with AlphaFold3, Boltz-2, Chai-1, and RosettaFold-3 across different MSA and template settings. Blue bars: close to groundtruth predictions, darker blue showing the lowest ligand RMSD models. In magenta, the structurally most deviating model considering both ligand RMSD but also local pocket environment. Protein backbone RMSDs remain low across models, whereas ligand RMSDs are low for orthosteric inhibitor predictions, but elevated experimental allosteric structure. **B** Relationship between model confidence and ligand placement accuracy. Scatter plot showing protein confidence (mean protein pLDDT) vs. ligand RMSD. **C** Scatter plot showing ligand confidence (mean ligand pLDDT) vs. ligand RMSD, illustrating the correlation between ligand-specific confidence and placement accuracy across all evaluated predictions.

Across the orthosteric kinase systems, ligand RMSDs were often low once the global fold was reproduced, but these values did not consistently coincide with correct recovery of local pocket geometry or state-defining residue conformations (Figure 4A). The effects of biased MSAs or templates on recovery of the correct conformational state were less consistent across kinases and GPCRs. We found that introducing a bias through MSAs or templates can improve local folding accuracy and ligand placement: When predicting the kinase type II inhibitor RN277 bound to LRRK2, all unbiased predictions failed to reproduce the correct binding mode, whereas introducing biased MSAs enabled recovery of the novel binding-pocket conformation.

However, in two conditions Boltz-2 with base settings yielded the most accurate structure (for ALK2-ATP and Adenosie A2A Receptor). This included the ATP example, for which a closely related ligand was present in the training data (ALK2 with AMPPNP in PDB 6UNQ). Although not showing the lowest ligand RMSD, AlphaFold3 and RosettaFold-3 predicted near-native structures for ALK2 bound to the type I inhibitor RK783 under base settings, with only modest improvement in RMSD upon addition of biased input information.

In summary, incorporating biased information into AlphaFold3, either through shallow MSAs or templates derived from state-filtered databases, results in the models structurally closest to the ground truth in four cases.

### Model confidence metrics do not report state fidelity

When identifying a metric to guide selection of the optimal ligand conformation in the absence of ground-truth structural information, we observed that protein pLDDT showed no association with pose accuracy (Pearson r = +0.04, Figure 4B), whereas ligand pLDDT correlated moderately (r = -0.46, Figure 4C) indicating that local ligand confidence, but not overall fold quality, is informative for co-folding predictions. This relationship suggests that the confidence metric may provide limited but informative guidance when evaluating ligand poses. This analysis could not be performed for RF3, as the ligand pLDDT value in its outputs is fixed at 1.0, preventing meaningful comparison.

### AlphaFold3 consistently delivers the most accurate cofolding predictions

From a method-centric perspective, AlphaFold3 produced the most reliable predictions across the majority of evaluated targets. Boltz-2 ranked second overall, generating the best-performing model in two cases but also the worst-performing model in two others, indicating greater variability. Chai-1 proved the least dependable, particularly for GPCR targets, where substantial variability was observed, most notably in ligand placement. RF3 generally occupied an intermediate position, displaying more consistent performance than Chai-1 but not reaching the accuracy achieved by AlphaFold3.

## Discussion

In this study, we evaluated state-of-the-art protein–ligand cofolding models on systems whose biological function depends on defined conformational states. Because these methods are primarily intended for applications with limited prior structural knowledge, we adopted a general, database-driven biasing strategy based on state annotations from GPCRdb (https://gpcrdb.org/)^6^ and KLIFS (https://klifs.net/)^7^ rather than target-specific manual curation. This design allowed us to probe how current cofolding frameworks behave under conditions that approximate realistic prospective use. Across the evaluated targets, accurate recovery of ligand-bound structures was achievable, but depended strongly on the interplay between model architecture and input biasing. In several cases, state-aware biasing enabled near-native predictions, whereas in others comparable accuracy was obtained without explicit biasing.

Because baseline templating failed to improve most predictions, template signals may need to be weighted more strongly to impact cofolding performance. This adjustment could be beneficial when ligands interact with well-established protein conformations. However, for conformationally novel or non-classical binding sites, excessive template weighting may bias models toward known inactive states, as seen in Boltz 2 with kinases.

The strong misfolding when omitting MSAs in RF3 and AF3 implies that MSA modifications influences AF3 and RF3 more strongly and would need to be applied with particular care to avoid destabilizing their predictions. At the same time, any custom MSA editing likely benefits from a more cautious approach if the goal is to preserve and possibly clarify evolutionary signal quality, which resonates with recent work on MSA denoising. ^29^ This observation is consistent with recent studies indicating that cofolding accuracy is dominated by strong structural priors, while shallow perturbations of evolutionary input alone often fail to reliably steer predictions.^20^

The kinase examples illustrate that state recovery remains a primary bottleneck uncoupled from correct ligand placement. Binding of RN277 in LRRK2 or ATP in ALK2 both require a defined orientation of a tyrosine, which was only resolved in a subset of predictions suggesting that cofolding models still depend strongly on training-set coverage and are influenced by the frequency of rare conformations within the templates too. For novel local pocket conformations, ligand placement was accurate, and biased inputs improved the local pocket geometries slightly but not consistently. The limitation of relying solely on filtered template databases becomes evident here: these datasets are enriched for canonical conformations (DFG-in/out), constraining cofolding models to familiar structural states. This constraint likely contributed to Boltz-2’s inability to reproduce the novel tyrosine conformation in LRRK2, even though its ligand placement remained accurate. A similar decoupling of ligand positioning from local backbone adaptation was reported by,^21^ who found that even removal of key binding residues did not substantially alter ligand positioning. This also supports the emerging view that generative structure predictors extrapolate only cautiously beyond known conformational space rather than fully sampling it.

Interestingly, for GPCRs we found that even when the ligand was placed reasonably, the active-state conformation was still consistently pulled toward the inactive state. That mismatch suggests that ligands exert only a modest influence on the global fold prediction and primarily shape the local environment of the binding pocket. This is helpful for interpreting binding interactions but limits the ability of specific ligands to steer the overall conformation in systems like GPCRs. However, suboptimal protein fold seems to effect the prediction of ligand conformations in a more drastic manner, as seen by the DOR example.

Active-state GPCRs appear to be particularly challenging for cofolding models, likely because activation involves substantial structural rearrangements that can differ between receptors. Intracellular loops, especially ICL3, are often unresolved in PDB structures due to their inherent flexibility. This lack of structural data may make accurate modeling more difficult. For instance, the 5-HT_2A_ receptor has an unusually long ICL3, and the absence of comparable templates for such extended loops could complicate full-length predictions even when the orthosteric ligand is correctly placed. The DOR was modeled with a G protein-biased agonist, ADL5859, which is a typ of ligand underrepresented in available co-crystallized structures. As a result, the intracellular conformation induced by this ligand may be uncommon and poorly sampled in template databases. Overall, variability in activation-related rearrangements, unresolved or unusually long intracellular loops, and limited structural coverage of biased ligand-bound states may contribute to the greater difficulty of modeling active-state GPCRs while ligand placement in the orthosteric site remains comparatively robust.

In our evaluation, per-residue pLDDT profiles were generally highest across the receptor core and decreased in flexible peripheral regions which we omitted in further analyses (Supplementary Information S2). Within the core, ICL3 emerges as one of the most challenging region to model accurately for GPCRs. This difficulty may stem from the models being trained on a mixture of active and inactive structures, creating ambiguity in this flexible intracellular loop, which undergoes the largest rearrangement during GPCR activation. Attempts to guide predictions through MSA editing or templating proved to be insufficient, especially for the active-state conformation, as there seems to be a preference for ICL3 conformations resembling the inactive state. In generell, models tend to predict the inactive-like ICL3 conformation almost perfectly, while their overall protein predictions for active or biased receptors reflects inactive-like or intermediate ICL3 structures. An exception in our evaluations was the DOR, where AF3 combined with a template and biased MSA correctly predicted ICL3. This suggests that, with additional weighting of these model biases, current methods may be able to better capture relevant protein conformations, ultimately improving the overall accuracy.

In this setup, custom templates were selected automatically based on the MSAs generated from the state annotated databases, and no additional criteria were applied beyond this sequence and state information. This meant that aspects such as ligand identity or functional context were not explicitly considered during template selection. In principle, the process could be made more deliberate by incorporating searches for templates with similar ligands, or, in cases involving biased GPCR agonists, by prioritizing structures that contain ligands with comparable signaling profiles. However, in the case of biased agonists, it remains unclear whether ligand bias is a broadly transferable feature, either across different GPCRs or even among ligands acting on the same receptor. Because of that uncertainty, any attempt to incorporate bias driven criteria into template selection would need to be approached cautiously. Still, it points to a possible way of broadening template choice beyond the purely sequence based strategy used here.

Furthermore, conformational states that are rare, mutation-dependent, or only accessible through dynamic reorganization are not recovered without explicit conditioning. In our study this was the case for allosteric ligands regulated by cryptic pocket opening^23^ and less conventional activation types such as biased agonists. Since they are markedly underrepresented in the structural data that most cofolding models were trained on, the models may not have a strong prior for how an allosteric pocket should be occupied or how its engagement reshapes the surrounding conformation. This limitation is consistent with recent analyses showing that cofolding models tend to overfit to well-represented protein–ligand interactions and struggle to generalize to uncommon binding sites or underrepresented conformational states, including allosteric pockets.^20^ In situations like this, using templates that contain a ligand bound in the relevant allosteric site could, at least in principle, help guide the model toward a more accurate arrangement of that region. This does not alter the overall workflow, but it highlights how template selection could be adjusted when the available structural landscape is sparse in ways that matter for the target system.

In conclusion, cofolding approaches appear promising due to their ease of use and adaptive fitting of the binding pocket to the ligand, which is lacking in many docking approaches. However, the overall reliability of the resulting protein conformations remains questionable, particularly with respect to state-specific accuracy. Incorporation of state-relevant, high-quality templates and multiple sequence alignments improves predictive performance. Moreover, careful ligand selection is crucial, as cofolding performs rather robustly for orthosteric ligands, it tends to fail for allosteric binders, highlighting a current limitation that must be considered when interpreting the results.

## Methods

### State specific MSA and template generation

For GPCRs, active and inactive state structures were retrieved from the PDB as mmCIF files, based on their state annotations in the GPCRdb (https://gpcrdb.org/).^6^ For kinases, the entire KLIFS database (https://klifs.net/)^7^ was downloaded, representing the most comprehensive collection of human kinases. PDB entries were sorted according to their conformational states, including DFG-out, DFG-inter, Active, C-Helix-out, and Salt-Bridge-out, using Kincore. ^30^ The selected PDB files were then converted to mmCIF format. Using Biopython,^31^ FASTA sequences corresponding to the preferred chain for each entry were generated. Multiple sequence alignments (MSAs) of each sub-database containing one conformational state were subsequently produced using MMseqs2^32^ saved as .a3m files using the `result2msa` command line option.

Custom templates were chosen by using the PDB structure files of the top four hits in each individual state-specific MSAs.

### Unbiased MSAs and Template generation

For Boltz-2 and Chai-1, MSAs were obtained via their MSA-server mode, which submits query sequences to the public ColabFold MMseqs2 server (https://api.colabfold.com/).^12,14,33^ As RosettaFold-3 does not provide a built in MSA generation command, the MSAs were generated separately with the standalone ColabFold Docker image in msa-only mode using default settings. AlphaFold3 generated MSAs with its internal data pipeline, using Jackhmmer and Nhmmer searches against the local AlphaFold3 sequence databases^12^ downloaded 17.05.2025. Among these methods, only Chai-1 offers a template-server mode. When enabled, template hits are retrieved from the ColabFold MMseqs2 server in the form of a `all_chain_templates.m8` file that is parsed by the Chai-1 template loader.

### Inference

For each method and condition, the general inference settings were kept similar using one seed (“0”) and five diffusion samples. Further inference settings were kept on default if not indicated.

#### AlphaFold3

Five setups were used in AlphaFold3 for each protein. Custom MSAs were first processed as AlphaFold3 requires a specific formatting. Therefore, the state-specific MMseqs2 A3M alignments were normalized with reformat.pl (HH-suite v3.3.0^34^) using the `-M first` option, which filters sequences so that all entries contain the same number of non-lowercase characters as the query sequence. The processed MSAs were then provided to AlphaFold3 within the JSON files by setting the `“unpairedMsaPath”`entry and providing an empty string to `“pairedMsa”`. In the *no MSA* conditions, the `“unpairedMsaPath”` was set to an empty string too, while the *base settings* condition lacks the JSON field entirely, resulting in AlphaFold3 automatically generating MSAs with the internal data pipeline.

In the conditions containing templates, the top 4 MSA hits were used as templates for structural templating. Therefore, a python script was created that extracts sequences from mmCIF files and aligns the query and template sequences using BioPython^31^ before calculating the queryIndices and templateIndices arrays for the JSON input file. AlphaFold3 does not intend to combine an unbiased MSA with biased templates within one run, which is why we only generated 5 conditions for alphaFold3. To generate five diffusion samples, we used the `--num_diffusion_samples=5` flag. All models were generated with AlphaFold3 v3.0.1.^12^

#### Boltz-2

Boltz-2^13,14^ models were generated using six different setups. The *unbiased* models were created using the base input, which included the protein sequence and ligand SMILES, with multiple sequence alignments (MSAs) generated through the `--use_msa_server` flag. The *no MSA* models were produced by explicitly disabling MSA usage, setting the MSA input to “empty”. For the *template* condition, templates were selected from the top four hits identified in the MSA of the annotated database. The *biased MSA* models used a manually curated A3M file as input, generated according to the procedure described in the MSA generation section. For each condition, diffusion samples were set to five via the `--diffusion_samples 5` flag during inference. All models were generated with Boltz version 2.2.0. ^13,14^

#### Chai-1

For Chai-1, state-specific A3M MSAs were converted to Parquet format using `chai-lab a3m-to-pqt` and the resulting directory was supplied via the `--msa-directory` option during inference. Templates were provided through M8 alignment files, which define the correspondence between the query sequence and full-length protein data bank entries retrieved by the model. Because the custom templates from KLIFs database consist only of the kinase domains, the subject start and end indices were set to the kinase domain boundaries (0-based). To generate five diffusion samples `--num-diffn-samples 5` was added to the command during inference. All models were generated with Chai-1 version 0.6.1

#### RosettaFold-3

For RosettaFold-3 biased MSAs, the previously described A3M files were adjusted with a python script that strips lowercase insertions, enforces a fixed post-processed length equal to the query, and rewrites all sequences in single-line format. The colabfold-MSA was generated with ColabFold. Except for the condition without MSA, the resulting A3M files were supplied to RF3 via absolute `msa_path` entries in the JSON files. As RosettaFold-3 defines templating different than the other methods, templates were not considered. To generate five diffusion samples the `--diffusion_batch_size=5` flag was used. All models were generated with RosettaFold-3 (commit b5c9685c7) using the model rf latest.pt downloaded on 08.10.2025. ^16^

### Analysis

The generated cofolding structures were aligned to their corresponding crystal structure using PyMOL and protein RMSD was calculated simultaniously. For GPCRs, alignment and protein RMSD calculations were restricted to the folded receptor core to exclude highly flexible terminal and loop regions. Subsequently, the aligned complexes were saved as a PDB file. Ligand RMSD was evaluated with MDAnalysis based on the aligned PDB structures. Per-residue pLDDT profiles were generated from the highest-mean-protein-pLDDT prediction of each method.

## Data availability

The ground truth structures are available through the RCSB protein data bank (https://www.rcsb.org/)^35^ under the accesion codes: GPCRs: 8ZMG,^26^ 8UGW,^27^ and 8Y45;^28^ kinases: 6UNQ,^25^ 9L04,^24^ 9DMI^22^ and 8TSD.^23^ The generated cofolding structure models and analysis scripts are available via zenodo (https://doi.org/10.5281/zenodo.19481962).

## Supporting Information Available

- Supplementary Information of “Assessing State-Specific Accuracy of Cofolding Models for kinases and GPCRs”
  - S1 Per-Residue pLDDT of Cofolding Kinase Predictions
  - S2 Per-Residue pLDDT of Cofolding GPCR Predictions

## Author contribution

L.O. and N.P.D contributed equally to this work. L.O. and N.P.D conceptualized the study, conducted the computational studies and analysis of the results. P.K. and G.W provided supervision of the study. L.O. and N.P.D wrote the original manuscript. P.K. and G.W reviewed and edited the manuscript. All authors have read and approved the manuscript.

## TOC Graphic

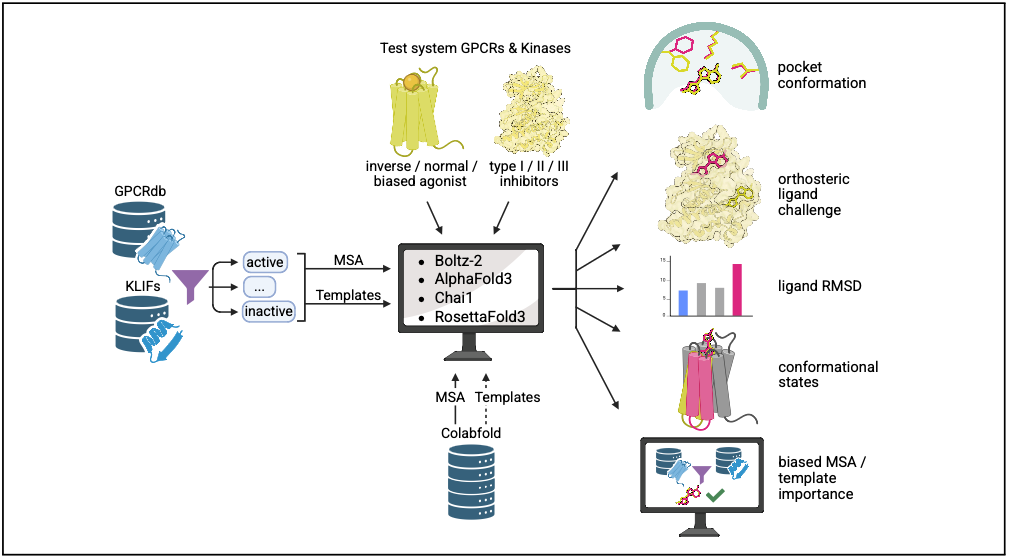

