## Supplemental Figures for "Assessing State-Specific Accuracy of Cofolding Models for Kinases and GPCRs"

#### Contents

**1 S1 Per-Residue pLDDT of Cofolding Kinase Predictions 2**

**2 S2 Per-Residue pLDDT of Cofolding GPCR Predictions 3**

### 1 S1 Per-Residue pLDDT of Cofolding Kinase Predictions

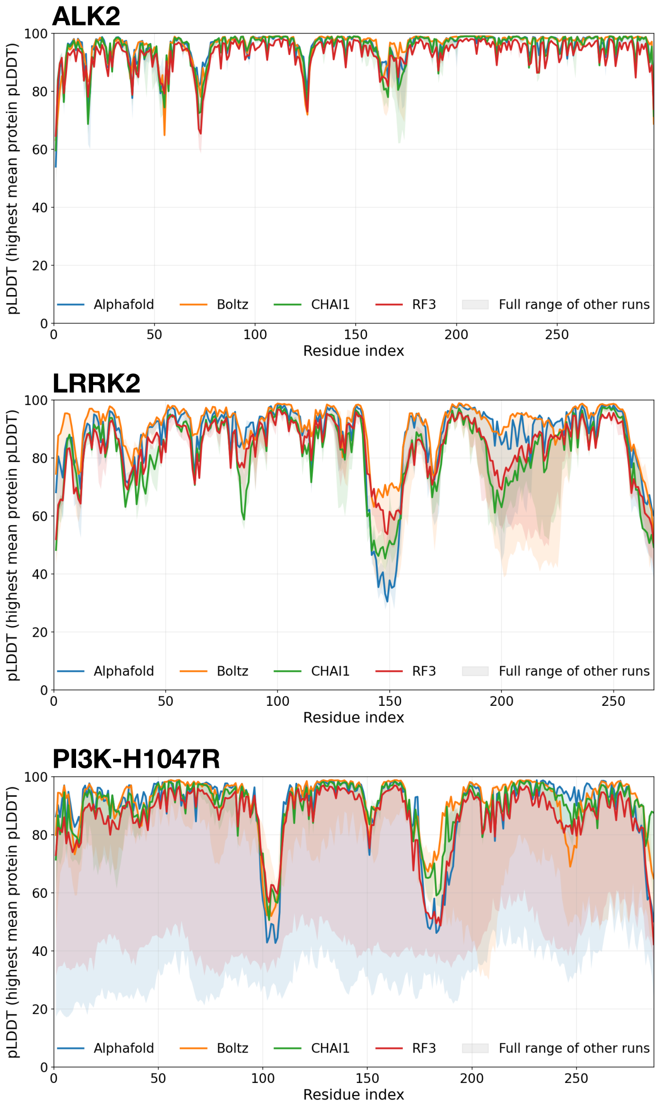


**Fig. 1** Per-residue pLDDT profiles for the best single kinase structure per method (colored lines), selected by highest mean protein pLDDT. The color-shaded region shows the full range (min–max) of pLDDT values across all other runs (besides the failed AlphafFold and RosettaFold no MSA / no Templates) for each residue position. Residue indices are shown on the x-axis; pLDDT scores (0–100) are on the y-axis.

### 2 S2 Per-Residue pLDDT of Cofolding GPCR Predictions

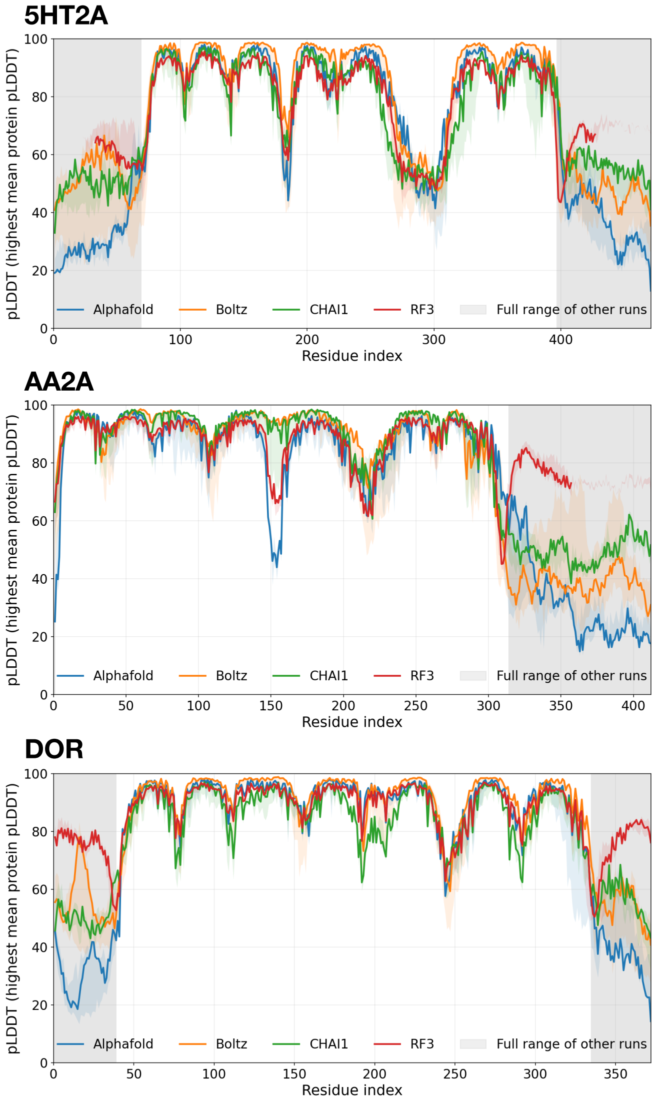


**Fig. 2** Per-residue pLDDT profiles for the best single GPCR structure per method (colored lines), selected by highest mean protein pLDDT. The color-shaded region shows the full range (min–max) of pLDDT values across all other runs (besides the failed AlphafFold and RosettaFold no MSA / no Templates) for each residue position. Residue indices are shown on the x-axis; pLDDT scores (0–100) are on the y-axis. The grey shaded region shows flexible parts of the GPCRs which were not used during alignment and plddt-comparison, to only compare the core GPCR besides N- or C-terminal loops.
